# Picophytoplankton dominate phytoplankton communities in the Elbe estuary in terms of cell counts

**DOI:** 10.1101/2024.05.31.596827

**Authors:** Nele Martens, Johanna Biederbick, C.-Elisa Schaum

## Abstract

(1) Picophytoplankton are important primary producers, but not always adequately recognized, e.g. due to methodological limitations.
(2) In this study, we combined flow cytometry and metabarcoding to investigate seasonal and spatial patterns of picophytoplankton abundance and community composition in the Elbe estuary.
(3) Picophytoplankton (mostly picoeukaryotes such as *Mychonastes* and *Minidiscus)* contributed on average 70 % (SD = 14 %) to the total phytoplankton counts. In the summer picocyanobacteria (e.g. *Synechococcus*) played a more significant role. The contributions of picophytoplankton to the total phytoplankton were particularly high from summer to winter as well as in the mid estuary. However, at salinities of around 10 PSU in the mixing area the proportion of picophytoplankton was comparably low (average 49 %, SD = 13 %).
(4) Our results indicate that picophytoplankton prevail in the Elbe estuary year-round with respect to cell counts. Picophytoplankton could occupy important niche positions to maintain primary production under extreme conditions where larger phytoplankton might struggle (e.g. at high or low temperature and high turbidity), and also benefit from high nutrient availability here. However, we did not find evidence that they played a particularly significant role at the salinity interface. Our study highlights the importance of including picophytoplankton when assessing estuarine phytoplankton as has been suggested for other ecosystems such as oceans.

## 1. Introduction

Picophytoplankton (< 2-3 µm) are important primary producers in aquatic ecosystems from oligotrophic to eutrophic habitats (Coello-Camba and Agustí, 2021; Moreira-Turcq et al., 2001; Purcell-Meyerink et al., 2017; Takasu et al., 2023; Zhang et al., 2015). These tiny organisms fulfill crucial ecological functions, e.g. as food for nauplii larvae and filter feeders (Bemal and Anil, 2019; Richard et al., 2022) and in carbon export (Basu and Mackey, 2018; Puigcorbé et al., 2015). The small size of picophytoplankton allows them to occupy specific ecological niches, for example due to the high surface to volume ratio which might facilitate the uptake of required nutrients, and slow sinking velocity that can keep them in the euphotic zone (Massana, 2011; Raven, 1998). Short generation times and high standing genetic variation give picophytoplankton a comparatively high evolutionary potential (e.g. (Barton et al., 2020; Benner et al., 2020; Schaum et al., 2016)). Picophytoplankton are more than likely to prevail in changing environments (see e.g. (Benner et al., 2020; Flombaum and Martiny, 2021a; Tan et al., 2022)). Some picophytoplankton have been shown to appear under extreme conditions e.g. at high or varying salinity, turbidity and temperature (Belkinova et al., 2021; Somogyi et al., 2022).

Extreme living conditions are common across ecosystems, including estuaries. Estuaries are the interfaces between the freshwater and marine world and characterized by gradients and tidal-induced variation of environmental forcing (e.g. salinity, turbidity) and picophytoplankton can be an important group here (Gaulke et al., 2010; Moreira-Turcq et al., 2001; Purcell-Meyerink et al., 2017; Sathicq et al., 2020). However, due to their small size picophytoplankton are still often not adequately recognized. This is largely due to difficulties in detecting and identifying these small-celled organisms with light microscopy, and because they cannot always be thoroughly preserved (Bergkemper and Weisse, 2018; Huo et al., 2020).

Here, we applied flow cytometry and metabarcoding (partially from (Martens et al., 2024b), in the context of new metabarcoding data) to (1) investigate spatial and seasonal patterns in picophytoplankton abundance and composition in the Elbe estuary, (2) identify dominant taxa and (3) assess under which conditions (with respect to abiotic factors, season, location) picophytoplankton and different players within (e.g. picocyanobacteria) might be particularly dominant.

## 2. Material and Methods

The Elbe estuary is located in the North of Germany, passing through the city of Hamburg, and enters the North Sea at Cuxhaven (fig. 1a). As one of Europe’s largest estuaries it is an important natural habitat and supplies the human population with essential ecosystem services (e.g. *via* port of Hamburg, recreation areas). The Elbe estuary has been experiencing intense anthropogenic pressure for centuries and further changes such as global warming or deepening of shipping channels might have additional impacts on the ecosystem functioning (see e.g. (Van Maren et al., 2015)). The tidal estuarine area is separated from the Elbe river by a weir at 586 km distance from the river source. Here, a total of 50 surface water samples were taken from seven stations along the Elbe estuary during different sampling campaigns (fig. 1a, supplementary tab. S1). Samples were taken from the upper water layers. Further details about sampling in the different sampling campaigns - e.g. sampling method and sample volume - are given in the supplementary data (tab. S1). 25 of the samples were taken around the city of Hamburg (approx. 623 - 633 km) and used as a seasonal dataset (fig. 3, fig. S4) and 29 samples from longitudinal sampling of six stations (609 - 713 km) covering three different seasons (spring and summer each 2021 and 2022 as well as winter 2022) were used as a spatial dataset (fig. 2a-c, fig. S2).

**Fig. 1:**
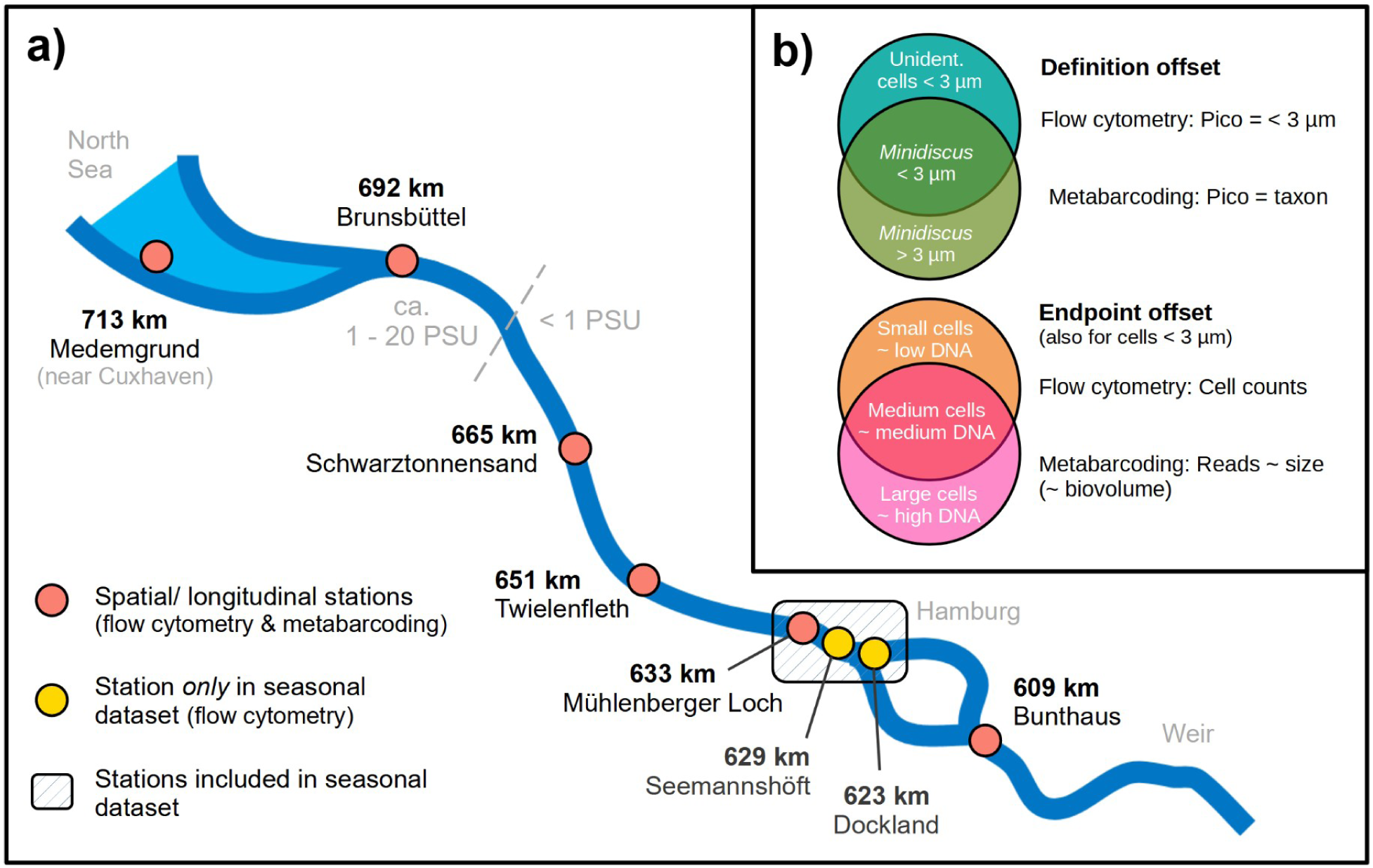
Study area (Elbe estuary) and sampling stations of the seasonal and spatial dataset (a) and schematic overview of the overlap and offset between the measured endpoint and definition of picophytoplankton (Pico) by the different methods (b). In (a) “km” metric indicates the approximate distance from the spring of the Elbe river in the Czech Republic. The discontinuous line shows the approximate border between freshwater and increased salinities during the sampling campaign. In (b) text within the circles are examples.

**Fig. 2:**
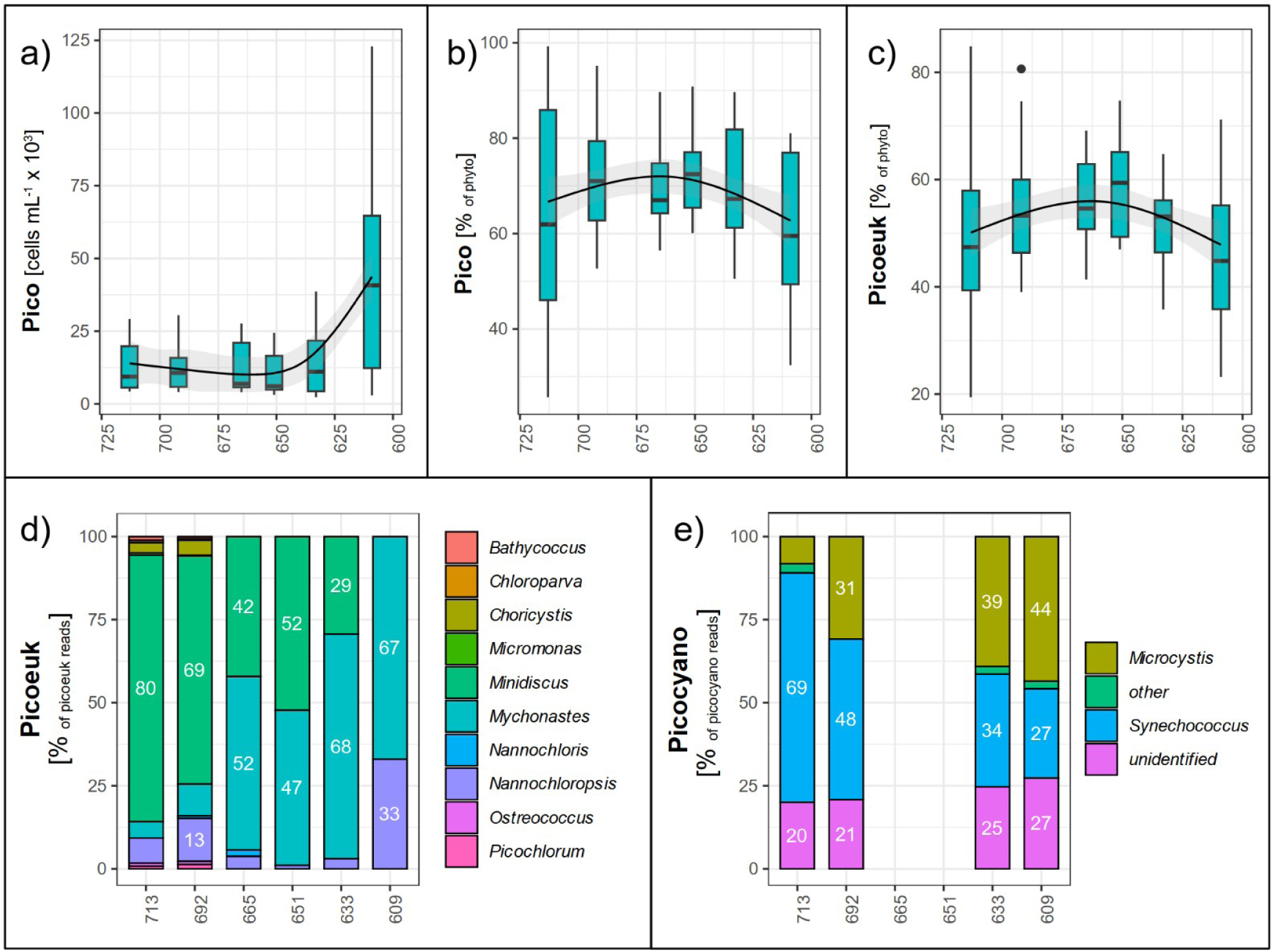
Spatial distribution of different picophytoplankton groups along different stations (stream km) across different seasons. Additional data, also separated by season, can be found in the supplementary material (fig. S2, fig. S3). Note that the number of samples included per season might differ, especially in d), and e) only shows data from summer 2021 due to lack of sufficient data from other seasons. In a) and e) station is used as a factor for clarity and labels are shown for values >= 10 %. Contributions of cyanobacteria to picophytoplankton can be obtained from c) (“picocyanobacteria = 1 - picoeukaryotes”). Regression lines were added with geom_smooth() from ggplot2 and the method “gam” (see also tab. S3). For clarity we use the following abbreviations: Pico = picophytoplankton, picoeuk = picoeukaryotes, phyto = phytoplankton. a) - c) refer to cell counts from flow cytometry, d) - e) to reads from metabarcoding. Data in d) is obtained from a former study (Martens et al., 2024b).

**Fig. 3:**
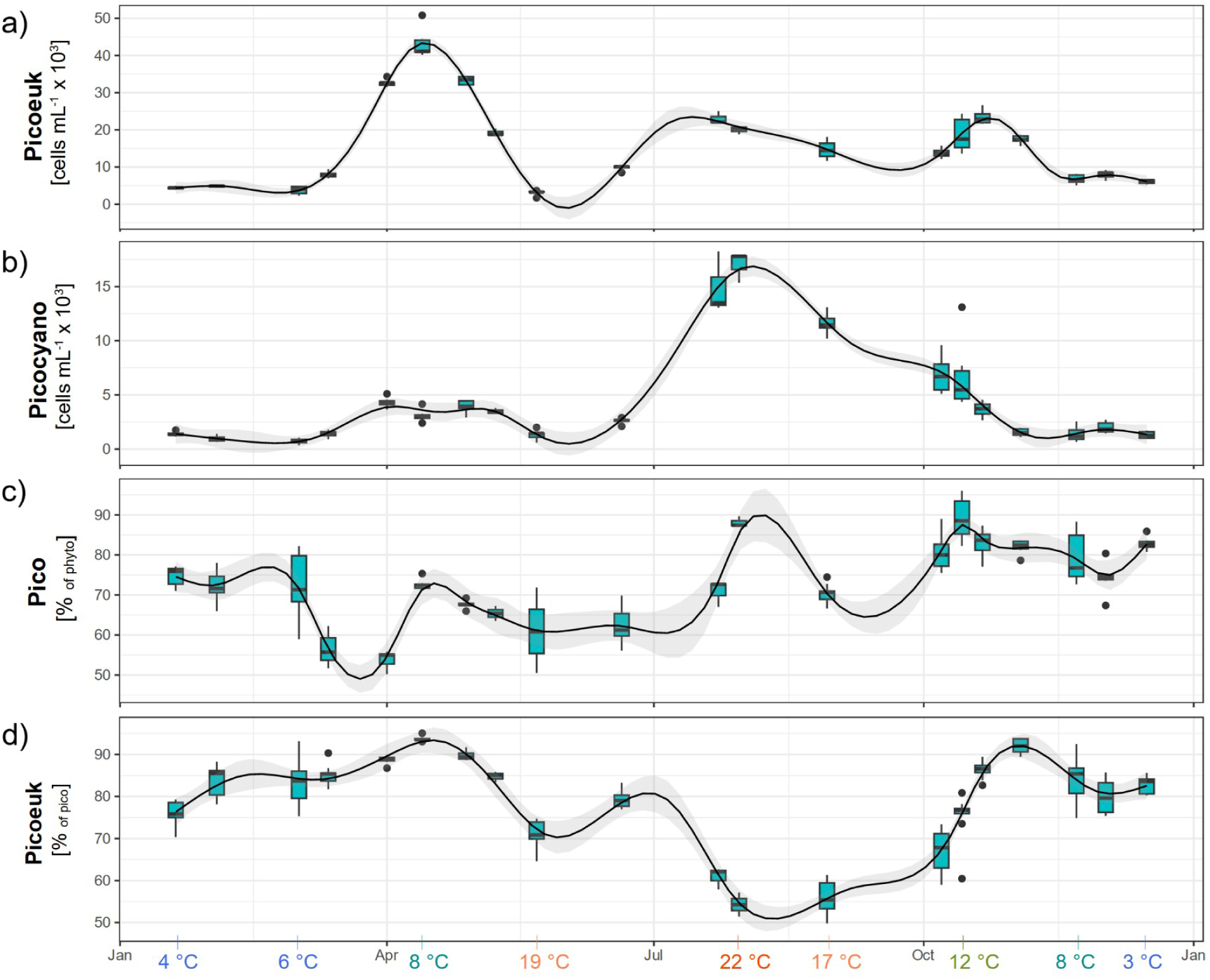
Seasonal distribution of different picophytoplankton groups in the area around Hamburg (approx. 623 - 633 km). Additional data can be found in the supplementary material (fig. S4). Horizontal scales show the sampling date independent of the year, i.e. day of the month. Data was merged when sampling was carried out < 5 days apart. Contributions of cyanobacteria to picophytoplankton can be obtained from the picoeukaryotes contribution in c) (“picocyanobacteria = 1 - picoeukaryotes”). Regression lines were added with geom_smooth() from ggplot2 and the method “gam” (see also tab. S3). For clarity we use the following abbreviations: Pico = picophytoplankton, picoeuk = eukaryotes, phyto = phytoplankton. On the bottom we show the temperatures at certain time points (see further details in fig. S6).

Of each sample, 3 to 5 technical replicates à 20 µL were analyzed using flow cytometry (BD accuri C6 plus) with a flow rate of 66 µl min^-1^ and regular cleaning and mixing between the samples. Phytoplankton cells could be distinguished from other suspended matter by their cytometric properties (e.g. fluorescence, size) which were also used to identify different groups of phytoplankton (see e.g. (Ning et al., 2021; Read et al., 2014; Thyssen et al., 2022) and supplementary material fig. S1). Picophytoplankton in the included samples from the Elbe estuary could be divided into two major groups: Picoeukaryotes and picocyanobacteria. Picocyanobacteria differed from picoeukaryotes in their fluorescence properties. This group had a higher phycocyanin- and lower chlorophyll-fluorescence (fig. S1). Notably, some larger cells might be excluded from our analysis due to detection limits and low sample volume. However, we know from former data (see e.g. (Martens et al., 2024b; NLWKN, 2023)) that taxa < 40 µm (e.g. *Stephanodiscus*, *Cyclotella*) were dominant in most seasons and areas of the Elbe estuary.

For the spatial dataset, 16S rRNA metabarcoding as well as 18S rRNA metabarcoding from another study (see further information in (Martens et al., 2024b)) were included to add information about picophytoplankton taxa in the Elbe estuary (fig. 2d-e, fig. S3). Samples for 16S rRNA sequencing were processed in the same way as shown for the 18S data (Martens et al., 2024b), however, reads were assigned using the BLAST database (carried out by biome-id Dres Barco & Knebelsberger GbR). In both datasets, we selected taxa that are in general considered picophytoplankton or can be < 3 µm (e.g. *Synechococcus*, *Choricystis*, *Mychonastes, Minidiscus*). We kept colony forming taxa that might appear solitary where single cells *can* be < 3 µm (e.g. *Microcystis*) - as well as unidentified cyanobacteria - in the dataset as they might add to the picocyanobacteria counts in the cytometry data. Note that the definition of picophytoplankton in the metabarcoding data is based on taxa identity and their *usual* size ranges, while in flow cytometry the definition is exclusively based on the *actual* cell size (< 3 µm) (fig. 1b). Furthermore, while in flow cytometry we detect abundance, metabarcoding results are rather correlated with biovolume. This is due to the size dependence of DNA copies per cell (Godhe et al., 2008) and as a result, larger (picophytoplankton) taxa might appear more dominant in metabarcoding compared to flow cytometric data without being more abundant in terms of cell counts. Consequently, what is included in “picophytoplankton” and how dominant it is can to some extent differ between the methods (see fig. 1b for further examples/ details). We also excluded data from samples with less than 100 picocyanobacteria, respectively picoeukaryotes reads in metabarcoding (fig. S3). The number of picophytoplankton reads per sample varied from 147 to 6108 (average 1733) in the 18S dataset and 113 to 5692 (average 2155) in the 16S dataset.

Data were processed in R (version 4.1.3), including the packages tidyverse (version 1.3.2), ggplot2 (version 3.4.0), lubridate (version 1.9.2), scales (version 1.2.1) and MuMIn (version 1.47.5). We also used LibreOffice Draw (version 7.1.2.2) for overview figures and addition of text notes. For spatial analyses, we obtained potentially interesting patterns from the figures showing cell counts and contributions of picophytoplankton groups along stations (fig. 2a-c, fig. S2) and then carried out an ANOVA aov() and Tukey test TukeyHSD() from the package stats (version 4.3.1) to assess whether the observed patterns were significant. To do so, we partially clustered different stations together, precisely the mid estuary (633 - 692 km) and the mid to lower estuary (633 - 713 km) and compared those with the residue stations, i.e. the uppermost (609 km) and lowermost (713 km) station. In the figures 2a-c, 3 and S4 with used GAMs for curve fitting with geom_smooth() from ggplot2 and the formula y ∼ s(x, bs = “cr”, k). The k value was determined based on the lowest AIC as obtained from uGamm() from the package MuMIn and AIC() from stats (see tab. S3). A spearman rank correlation with the function rcorr() from the package Hmisc() (version 5.1-0) was applied to draw conclusions about the relationship of picophytoplankton groups with abiotic parameters obtained from the samples (precisely water temperature, salinity, turbidity, PO_4_ and NO_3_; see also supplementary data fig. S5-S7 and tab. S4) (fig. 4). Those data were provided by Helmholtz-Zentrum hereon and obtained from the FGG database (FGG Elbe, 2024).

**Fig. 4:**
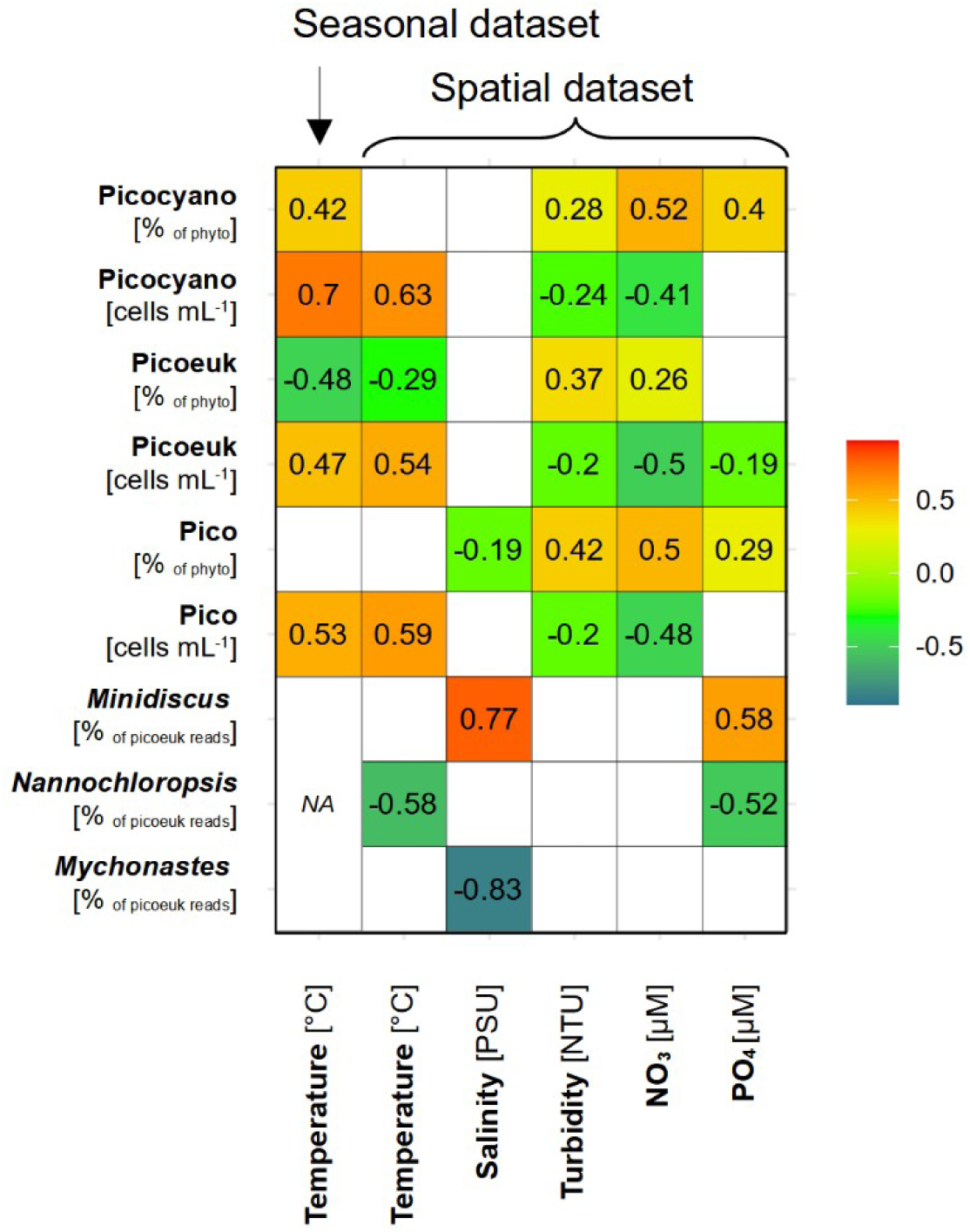
Correlation of different picophytoplankton groups with abiotic conditions in the spatial and seasonal dataset. Numbers and color scheme show the correlation coefficient r calculated with spearman rank correlation for p ≤ 0.05 (see also tab. S4). Salinity, turbidity, NO_3_ and PO_4_ are shown for the spatial dataset only, as these parameters did either not vary much or were not obtained in the seasonal dataset. Temperature is shown for both datasets separately. 16S data was not included here due to the low number of data points (see methods). For clarity we used the abbreviations picoeuk = picoeukaryotes, picocyano = picocyanobacteria and phyto = phytoplankton.

## 3. Results

Across seasons and methods, flow cytometry detected between 2.3 x 10^3^ and 123 x 10^3^ picophytoplankton cells mL^-1^ in the samples from the different stations and seasons from the Elbe estuary. On average 70 % (SD = 14 %) and up to 99 % of the detected phytoplankton cells per sample were < 3 µm. Picoeukaryotes were by far the most dominant group with an average contribution of 77 % (SD = 11 %) to the picophytoplankton cell counts, while picocyanobacteria played a role in summer (up to 53 %).

Across seasons, picophytoplankton, picoeukaryotes and picocyanobacteria were overall significantly more abundant at the uppermost station (609 km) than in the area further downstream (633 - 713 km) (fig. 2a, ANOVA/ Tukey: p = 6.0 x 10^-11^, p = 2.1 x 10^-10^ and p = 3.7 x 10^-6^, respectively; see also tab. S2). Contributions of picoeukaryotes to the phytoplankton cell counts were significantly higher in the mid estuary (633 - 692 km) compared to the upper station (609 km) (fig. 2c, ANOVA/ Tukey: p = 0.009; tab. S2), while they showed no significant differences between the mid and lower, as well as the upper and lower stations (tab. S2). As the picophytoplankton fraction was largely represented by picoeukaryotes, those patterns hold for the contributions of picophytoplankton to the phytoplankton cell counts as a whole (fig. 2b, ANOVA/ Tukey: p = 0.031 for comparison of the mid (633 - 692 km) and upper area (609 km); tab. S2). In contrast, contributions of picocyanobacteria to the picophytoplankton did not express a distinct pattern along space across season (fig. S2f).

*Minidiscus* and *Mychonastes* were the most dominant picoeukaryote taxa across seasons based on 18S rRNA reads (fig. 2d). Therein, *Mychonastes* was more dominant in the upper to mid reaches of the estuary (approx. 609 – 665 km), and *Minidiscus* in the mid to lower area (approx. 651 – 713 km). *Nannochloropsis* was prominent at 609 km in early May (spring 2021) and at 692 to 713 km in February (winter 2022) (fig. S3a). Here *Choricystis* also played a role (contributions up to approx. 20 %). Other picoeukaryotes such as *Bathycoccus* and *Picochlorum* were minor contributors to the 18S picophytoplankton reads. Results from 16S sequencing (fig. 2e) show that *Synechococcus* and *Microcystis* might be the most relevant contributors to picocyanobacteria in summer 2021, where picocyanobacteria were particularly dominant (up to 53 % of the picophytoplankton cells; see also fig. S2f). Here *Microcystis* was more dominant at the upper stations (609 - 633 km) and *Synechococcus* at the lower stations (692 - 713 km). Notably there is some degree of uncertainty to what extent *Microcystis* would fall into the size range of picophytoplankton, due to colony formation and cell size. It is likely that *Synechococcus* reached significantly higher proportions among the cells < 3 µm than suggested in figure 2e. Minor contributors to the picocyanobacteria reads were e.g. *Prochlorococcus* and *Cyanobium* (“other” in fig. 2e).

In the seasonal dataset from downstream of the city of Hamburg (623 - 633 km), the abundances and contributions of the different picophytoplankton groups expressed distinct patterns along the sampling dates. The complexity is reflected in the high k values (15 - 20) of the fitted GAMs (fig. 2, fig. S4, tab. S3). Picoeukaryotes expressed three seasonal peaks in spring, summer and fall, where the spring peak was most pronounced (fig. 3a). Picocyanobacteria reached their maximum abundance around July to August with high abundances extending into October (fig. 3b). Picophytoplankton contributions to the total phytoplankton were highest in a single sample from the temperature peak in summer (fig. 3c), largely due to picocyanobacteria (fig. 3b+d), and across different samples in fall, which is due to low abundance of larger-celled taxa combined with the fall peak of picoeukaryotes and the remains of the fading summer bloom of picocyanobacteria (fig. 3a-b, fig. 3d). In contrast, picophytoplankton were less dominant within the phytoplankton communities in spring (fig. 3c) due to taxa > 3 µm blooming in parallel. Seasonal effects could also be observed in the spatial dataset as longitudinal data was obtained from different seasons (winter, spring and summer). Here, highest absolute abundances of picophytoplankton were observed in summer 2022 (fig. S2a). Contributions of picophytoplankton to the total phytoplankton counts were overall highest in summer 2021 and in winter 2022 mostly due to picocyanobacteria as well as picoeukaryotes and low abundance of larger-celled phytoplankton, respectively (fig. S2d-f).

Abiotic conditions varied along seasons and stations (see also supplementary fig. S5-S6). Temperature was highest in July (up to 23 °C in the spatial dataset at 665 km) and low in winter (down to 3 °C). Turbidity, salinity, NO_3_ and PO_4_ expressed spatial and seasonal patterns (fig. S5). Salinity was enhanced at the lowermost stations (692 - 713 km) and highest in summer 2022 and spring of both years (fig. S5a). Compared to other seasons, turbidity was enhanced in winter 2022 and spring 2021 (fig. S5b), NO _3_ in winter 2022 and summer 2021 (fig. S5c) and PO_4_ in summer 2021 (fig. S5d). Note that turbidity was also enhanced in fall 2021 (fig. S7; (FGG Elbe, 2024)) where we did not obtain longitudinal samples, but seasonal samples from a fixed area (623 - 633 km). Turbidity, PO_4_ and NO_3_ concentrations were enhanced downstream of 609 km (fig. S5b-d).

Picophytoplankton abundance was positively correlated with temperature (fig. 4). This was a result of high abundance of picocyanobacteria in summer 2021 (fig. 3b, fig. S2c, fig. S6), high abundance of picoeukaryotes in summer 2022 (fig. S2b) and low abundances of both groups in winter (fig. S2b-c). Relative contributions of picoeukaryotes to the (pico-)phytoplankton were negatively correlated with temperature (fig. 4). Overall this pattern arises from relatively high picoeukaryotes contributions to the phytoplankton in fall and winter (fig. S2e, fig. S4b) where phytoplankton abundance was generally low, and enhanced picocyanobacteria contributions to the picophytoplankton in summer (fig. 3d, fig. S2g). Picophytoplankton contributions to the phytoplankton cell counts were negatively correlated with salinity (fig. 4), largely due to low contributions at around 10 PSU at 713 km in spring 2021 and 2022 (fig. S2d, fig. S5a). Cell counts of picophytoplankton were negatively correlated with turbidity and NO_3_ (fig. 4) due to their high absolute abundance at 609 km - especially in summer - where turbidity and NO _3_ concentrations were rather low (fig. S2a, fig. S5b-c). In contrast, relative contributions of picophytoplankton to the phytoplankton were positively correlated with these parameters and additionally with PO_4_ (fig. 4). This relationship with PO_4_, NO_3_ and turbidity is affected by the higher proportions of small cells in the mid to lower estuary compared to 609 km, where these parameters achieved overall higher values and by the seasonal dominance of picocyanobacteria in summer 2021 (at high PO_4_) and picoeukaryotes in winter (at high turbidity and NO_3_) (fig. S2d-f, fig. S5b-d). Due to enhanced contributions to the picoeukaryotes reads from winter to spring compared to the other seasons (fig. S3a), *Nannochloropsis* was negatively correlated with temperature (fig. 4). The negative relationship with PO_4_ can be mainly explained by the high contributions of this taxon at 609 km in spring 2021, where PO_4_ was particularly low (precisely below detection limit; fig. S5d). *Mychonastes* was clearly associated with the freshwater reaches of the estuary (fig. 2d, fig. S3a), resulting in negative correlation with salinity (fig. 4, fig. S5a). In contrast, *Minidiscus* was more dominant further downstream and hence associated with higher salinity and higher PO_4_ values (fig. 4, fig. 2d, fig. S3a, fig. S5a+d).

## 4. Discussion

We used flow cytometry to quantify picophytoplankton along the Elbe estuary and across seasons, and combined the results with composition data obtained from metabarcoding. Our results indicate that picophytoplankton - and therein picoeukaryotes - were the dominant groups of phytoplankton in the Elbe estuary with respect to abundance in the vast majority of the samples. Notably, different picoeukaryote taxa (precisely *Minidiscus* and *Mychonastes*) could each contribute up to 17 % to the eukaryotic phytoplankton reads, implying that this group was also relevant in terms of biovolume (see also (Martens et al., 2024b) and fig. 1b). Considering their ubiquitous appearance throughout water bodies around the world (Coello-Camba and Agustí, 2021; Purcell-Meyerink et al., 2017; Sathicq et al., 2020; Takasu et al., 2023), it is not surprising that picophytoplankton also play an important role in the Elbe estuary, even though empirical evidence has so far been scarce for this ecosystem.

Picophytoplankton have been found to be important at extreme and highly variable salinities, e.g. in hypersaline lakes and in the Black Sea (Belkinova et al., 2021; Somogyi et al., 2022) and at intermediate salinities (e.g. 5 - 10 PSU) in estuaries (Wetz et al., 2011), which are somewhat extreme for both freshwater and saltwater inhabitants. However, our data so far imply that picophytoplankton were overall more abundant and dominant at freshwater and rather high salinity (approx. 20 PSU). Nevertheless, some picophytoplankton taxa, such as certain genotypes of *Minidiscus* (see also (Martens et al., 2024b)) as well as *Ostreococcus*, *Bathycoccus* and *Picochlorum*, which were particularly associated with intermediate salinities (approx. 1 - 10 PSU), have been associated with brackish habitats (e.g. in (Hu et al., 2016; Tragin and Vaulot, 2019)) and high salinity tolerances (Foflonker et al., 2016; Somogyi et al., 2022) before. Those groups might fulfill significant ecological functions at the salinity interface, e.g. as primary producers and as food items for higher trophic levels, the latter regardless of whether they are in particularly good condition.

Distribution patterns in the contributions of picophytoplankton to the total phytoplankton cells - and the relationship of those contributions with abiotic parameters - indicate that picophytoplankton play a major role under extreme environmental conditions with respect to temperature and turbidity. Picocyanobacteria were associated with high water temperatures in summer (e.g. up to 22 °C; see also temperatures along seasons in fig. S6), as observed for picocyanobacteria in other studies (Alegria Zufia et al., 2021; Murrell and Lores, 2004). This group may become more dominant in the Elbe estuary under global warming (see also (Flombaum and Martiny, 2021b)), which might affect food webs due to the relatively low nutritional value and possible toxicity of cyanobacteria (Ger et al., 2016; Sim et al., 2023). Picoeukaryotes were most abundant in spring, but their relative contributions to the phytoplankton communities were overall highest in fall and winter, as well as in the middle reaches of the estuary, i.e. at a combination of low temperature and low light availability (due to turbidity and sunlight). While further research is needed to disentangle the effects of temperature and turbidity, picoeukaryotes have been associated with the colder seasons in other studies (Alegria Zufia et al., 2021; Vörös et al., 2009) and a positive relationship between turbidity and picophytoplankton as a whole has been observed before. Those studies argue that picophytoplankton might have specific strategies in light harvesting (Coe et al., 2021; Malinsky-Rushansky, 2002; Somogyi et al., 2022, 2017; Soulier et al., 2022). Additionally, we found that picoeukaryotes from the Elbe estuary were particularly skilled in utilizing organic compounds (Martens et al., 2024a) which might give them advantages over other phytoplankton groups. The positive relationship of the contributions of different picophytoplankton groups with nutrients (NO_3_ and PO_4_) furthermore imply that those groups benefit from high nutrient availability, e.g. in areas with otherall low phytoplankton concentrations such as the mid to lower estuary (633 - 713 km).

Phytoplankton concentrations are known to decline along the Elbe estuary, especially through the city of Hamburg. This is partially explained by local grazing effects (Schöl et al., 2009) but may also be affected by e.g. sinking in the current-calmed harbor basins (Wolfstein, 1996). Picophytoplankton abundance followed this pattern in our study. However, due to their small size, they might be less affected by those factors than larger-sized taxa, which can beyond the aforementioned ones contribute to the elevated contributions in the mid estuary compared to the station upstream of Hamburg (609 km). For instance, small picophytoplankton cells have a reduced sinking velocity compared to larger-celled phytoplankton. Moreover, while one of the key zooplankton taxa - *Eurytemora* (Schöl et al., 2009) - may utilize picophytoplankton e.g. as part of aggregates (Modéran et al., 2012; Wilson and Steinberg, 2010) - they likely prefer to consume larger-celled phytoplankton, and hence, picophytoplankton might be removed less rapidly by grazing.

Methodological limitations - such as an underrepresentation of samples with intermediate to higher salinities - might have affected some of our interpretations, however, consistent findings across a high number of samples included, e.g. with respect to the picophytoplankton dominance in terms of cell counts, make it inevitable to conclude that picophytoplankton play a key role in the Elbe estuary. Their high contributions under extreme conditions - e.g. high temperatures and low light availability - implies that they occupy ecological niches to maintain primary production where larger phytoplankton might struggle, and their small nature might protect them from rapidly being removed from the water body. Our results emphasize the importance to include the so far underrated group of picophytoplankton in (estuarine) research and provide insights into the comparability of techniques (e.g. flow cytometry, metabarcoding) for detecting (pico)phytoplankton communities.

## Acknowledgements

We would like to thank the captain and crew of the research vessel Ludwig Prandtl and the fishing vessel Ostetal as well as Raphael Koll and Elena Hauten for their contributions to obtain samples, Vanessa Russnak, Leon Schmidth and Kirstin Dähnke (Helmholtz-Zentrum hereon) for providing environmental data, as well as Stefanie Schnell and Luisa Listmann for the technical support in the laboratory and Inga Hense and Hans-Peter Grossart for conceptual support.

## Conflict of Interest Statement

We declare we have no competing interests

## Supplementary material

**Tab. S1:** Metadata.

| Station (km) | Date | Included in dataset(s): | Technical replicates flow cytometry | Type of sampling | Cruise number | Sampling method | Water sample volume * | Storage Times (cold) ** |
| --- | --- | --- | --- | --- | --- | --- | --- | --- |
| 609 | 2022-06-21 | spatial | 5 | Prandtl (Ship) | LP220613 | Ferrybox | 500 mL | ca. 14 - 22 h |
| 633 | 2022-06-21 | spatial + seasonal | 5 | Prandtl (Ship) | LP220613 | Ferrybox | 500 mL | ca. 14 - 22 h |
| 651 | 2022-06-21 | spatial | 5 | Prandtl (Ship) | LP220613 | Ferrybox | 500 mL | ca. 14 - 22 h |
| 665 | 2022-06-21 | spatial | 5 | Prandtl (Ship) | LP220613 | Ferrybox | 500 mL | ca. 14 - 22 h |
| 692 | 2022-06-21 | spatial | 5 | Prandtl (Ship) | LP220613 | Ferrybox | 500 mL | ca. 14 - 22 h |
| 713 | 2022-06-21 | spatial | 5 | Prandtl (Ship) | LP220613 | Ferrybox | 500 mL | ca. 14 - 22 h |
| 609 | 2022-05-22 | spatial | 5 | Prandtl (Ship) | LP220522 | Ferrybox | 500 mL | ca. 7 - 13 h |
| 633 | 2022-05-22 | spatial + seasonal | 5 | Prandtl (Ship) | LP220522 | Ferrybox | 500 mL | ca. 7 - 13 h |
| 651 | 2022-05-22 | spatial | 5 | Prandtl (Ship) | LP220522 | Ferrybox | 500 mL | ca. 7 - 13 h |
| 665 | 2022-05-22 | spatial | 5 | Prandtl (Ship) | LP220522 | Ferrybox | 500 mL | ca. 7 - 13 h |
| 692 | 2022-05-22 | spatial | 5 | Prandtl (Ship) | LP220522 | Ferrybox | 500 mL | ca. 7 - 13 h |
| 713 | 2022-05-22 | spatial | 5 | Prandtl (Ship) | LP220522 | Ferrybox | 500 mL | ca. 7 - 13 h |
| 629 | 2022-04-28 | seasonal | 5 | Longterm (Pier) | NA | Bucket/ tube | 15 mL | 7 h |
| 629 | 2022-04-13 | seasonal | 5 | Longterm (Pier) | NA | Bucket/ tube | 15 mL | 6 h |
| 629 | 2022-04-01 | seasonal | 5 | Longterm (Pier) | NA | Bucket/ tube | 15 mL | 4 h |
| 633 | 2022-03-12 | seasonal | 10 ( 5 each) | Ostetal (Ship) | NA | Bucket/ tube | 2 x 15 mL (ebb & flood) | ca. 1 d |
| 629 | 2022-03-02 | seasonal | 5 | Longterm (Pier) | NA | Bucket/ tube | 15 mL | 28 h |
| 609 | 2022-02-28 | spatial | 5 | Prandtl (Ship) | LP220228 | Ferrybox | 500 mL | ca. 4 - 13 h |
| 633 | 2022-02-28 | spatial + seasonal | 5 | Prandtl (Ship) | LP220228 | Ferrybox | 500 mL | ca. 4 - 13 h |
| 651 | 2022-02-28 | spatial | 5 | Prandtl (Ship) | LP220228 | Ferrybox | 500 mL | ca. 4 - 13 h |
| 665 | 2022-02-28 | spatial | 5 | Prandtl (Ship) | LP220228 | Ferrybox | 500 mL | ca. 4 - 13 h |
| 692 | 2022-02-28 | spatial | 5 | Prandtl (Ship) | LP220228 | Ferrybox | 500 mL | ca. 4 - 13 h |
| 713 | 2022-02-28 | spatial | 5 | Prandtl (Ship) | LP220228 | Ferrybox | 500 mL | ca. 4 - 13 h |
| 629 | 2022-02-03 | seasonal | 5 | Longterm (Pier) | NA | Bucket/ tube | 15 mL | 5 h |
| 629 | 2022-01-20 | seasonal | 5 | Longterm (Pier) | NA | Bucket/ tube | 15 mL | 5 h |
| 629 | 2021-12-16 | seasonal | 5 | Longterm (Pier) | NA | Bucket/ tube | 15 mL | 30 h |
| 629 | 2021-12-02 | seasonal | 5 | Longterm (Pier) | NA | Bucket/ tube | 15 mL | 9 h |
| 629 | 2021-11-22 | seasonal | 5 | Prandtl<br>substitute (Pier) | NA | Bucket/ tube | 15 mL | 3 h |
| 629 | 2021-11-18 | seasonal | 5 | Longterm (Pier) | NA | Bucket/ tube | 15 mL | 7 h |
| 629 | 2021-11-03 | seasonal | 5 | Longterm (Pier) | NA | Bucket/ tube | 15 mL | 27 h |
| 623 | 2021-10-21 | seasonal | 5 | Dockland (Pier) | none | Bucket/ tube | 15 mL | < 1 h |
| 629 | 2021-10-21 | seasonal | 5 | Longterm (Pier) | NA | Bucket/ tube | 15 mL | 3 h |
| 623 | 2021-10-14 | seasonal | 5 | Dockland (Pier) | none | Bucket/ tube | 15 mL | < 1 h |
| 623 | 2021-10-13 | seasonal | 5 | Dockland (Pier) | none | Bucket/ tube | 15 mL | < 1 h |
| 623 | 2021-10-06 | seasonal | 5 | Dockland (Pier) | none | Bucket/ tube | 15 mL | < 1 h |
| 629 | 2021-10-06 | seasonal | 5 | Longterm (Pier) | NA | Bucket/ tube | 15 mL | 26 h |
| 633 | 2021-08-29 | seasonal | 18 (3 each) | Ostetal (Ship) | NA | Bucket/ tube | 6 x 15 mL<br>(3x ebb, 3x flood) | up to 14 h |
| 633 | 2021-07-30 | spatial + seasonal | 3 | Prandtl (Ship) | LP210725 | Ferrybox | 2 L | 13 h |
| 665 | 2021-07-30 | spatial | 3 | Prandtl (Ship) | LP210725 | Ferrybox | 2 L | 10 h |
| 692 | 2021-07-30 | spatial | 3 | Prandtl (Ship) | LP210725 | Ferrybox | 2 L | 8 h |
| 713 | 2021-07-30 | spatial | 3 | Prandtl (Ship) | LP210725 | Ferrybox | 2 L | 6 h |
| 609 | 2021-07-29 | spatial | 3 | Prandtl (Ship) | LP210725 | Ferrybox | 2 L | 10 h |
| 623 | 2021-07-23 | seasonal | 3 | Dockland (Pier) | NA | Bucket/ tube | 15 mL | < 1 h |
| 692 | 2021-05-08 | spatial | 3 | Prandtl (Ship) | LP210503 | Ferrybox | 2 L | ca. 11 h |
| 713 | 2021-05-08 | spatial | 3 | Prandtl (Ship) | LP210503 | Ferrybox | 2 L | ca. 9 h |
| 609 | 2021-05-07 | spatial | 3 | Prandtl (Ship) | LP210503 | Ferrybox | 2 L | ca. 36 h |
| 633 | 2021-05-07 | spatial + seasonal | 3 | Prandtl (Ship) | LP210503 | Ferrybox | 2 L | ca. 34 h |
| 651 | 2021-05-07 | spatial | 3 | Prandtl (Ship) | LP210503 | Ferrybox | 2 L | ca. 34 h |
| 665 | 2021-05-07 | spatial | 3 | Prandtl (Ship) | LP210503 | Ferrybox | 2 L | ca. 32 h |

\*the sample volume describes the sample taken from the water and here refers to flow cytometry. These samples are then mixed before an aliquot of 20 µL per replicate is used for the flow cytometric measurements. Sample volumes for metabarcoding are shown in the supplementary data of Martens et al. 2024 (https://doi.org/10.1016/j.envres.2024.119126). In metabarcoding, the whole sample is concentrated and processed, but only an aliquot of the DNA extract is used for the actual sequencing (see methods description in aforementioned study).
\*\*times samples were stored in the fridge or under comparable conditions prior to flow cytometric. Storage times for metabarcoding samples (times prior to freezing) can be found in the supplementary material of Martens et al. 2024 (https://doi.org/10.1016/j.envres.2024.119126), but are often similar to the storage times of flow cytometric samples.

**Tab. S2:** Results from ANOVA and Tukey test for assessment of spatial patterns (spatial dataset)

| Picophytoplankton group | Compared areas | p | F | Direction |
| --- | --- | --- | --- | --- |
| Picophytoplankton cells | Upper (609 km) vs.<br>mid to lower<br>(633 - 713 km) | $6.0 \times 10^{-11}$ | 51.6 | <u>609 km > other</u> |
| Picoeukaryotes cells | | $2.1 \times 10^{-10}$ | 48.2 | |
| Picocyanobacteria cells | | $3.7 \times 10^{-6}$ | 23.5 | |
| Picophytoplankton %<br>of phytoplankton | Upper (609 km) vs.<br>mid (633-692 km) | 0.009 | 5.26 | <u>mid > upper</u> |
|  | Upper (609 km) vs.<br>lower (713 km) | 0.610 |  | not relevant |
|  | Mid (633-692 km) vs.<br>lower (713 km) | 0.171 |  | not relevant |
| Picoeukaryotes %<br>of phytoplankton | Upper (609 km) vs.<br>mid (633-692 km) | 0.031 | 4.42 | <u>mid > upper</u> |
|  | Upper (609 km) vs.<br>lower (713 km) | 0.891 |  | not relevant |
|  | Mid (633-692 km) vs.<br>lower (713 km) | 0.118 |  | not relevant |

**Tab. S3:** GAM fitting. (for the spatial data (fig. 2), we tested all possible k values (3 - 6), for seasonal data (fig. 3, S4) we tested k values from 10 - 20. In all cases we selected the k with the lowest AIC for in the figures). For clarity, only a selection of the tested k values and their AIC are shown here.

| Figure | k | AIC | Comment |
| --- | --- | --- | --- |
| 2a (Picophytoplankton cell counts ~ station) | 4 | 1096.92 |  |
|  | **5** | **1096.86** | Chosen for figure |
|  | 6 | 1097.1 |  |
| 2b (Picophytoplankton as % of phytoplankton ~ station) | **3** | **1009.9** | Chosen for figure |
|  | 4 | 1010.3 |  |
| 2c (Picoeukaryotes as % of phytoplankton ~ station) | **3** | **969.4** | Chosen for figure |
|  | 4 | 970.0 |  |
| 3a (Picoeukaryotes cell counts ~ date) | 10 | 784.2 |  |
|  | **17** | **651.7** | Chosen for figure |
|  | 20 | 655.6 |  |
| 3b (Picocyanobacteria cell counts ~ date) | 10 | 509.3 |  |
|  | **15** | **470.1** | Chosen for figure |
|  | 20 | 471.4 |  |
| 3c (Picophytoplankton as % of phytoplankton ~ date) | 10 | 922.7 |  |
|  | 15 | 906.8 |  |
|  | **20** | **878.8** | Chosen for figure |
| 3d (Picoeukaryotes as % of picophytoplankton ~ date) | 10 | 881.5 |  |
|  | **15** | **819.9** | Chosen for figure |
|  | 20 | 820.8 |  |
| S4a (Picophytoplankton cell counts ~ date) | 10 | 787.5 |  |
|  | **15** | **710.2** | Chosen for figure |
|  | 20 | 713.6 |  |
| S4b (Picoeukaryotes as % of phytoplankton ~ date) | 10 | 933.4 |  |
|  | 15 | 893.8 |  |
|  | **20** | **880.8** | Chosen for figure |
| S4c (Picocyanobacteria as % of phytoplankton ~ date) | 10 | 841.2 |  |
|  | 15 | 784.9 |  |
|  | **20** | **781.1** | Chosen for figure |

**Tab. S4:** r and p values from spearman rank correlation. (for abbreviations and units see fig. 4)

| **Var1** | **Var2** | **p** | **r** |
| --- | --- | --- | --- |
| Temp_spatial | *Mychonastes* | 0.2698 | 0.27 |
| Salinity | *Mychonastes* | < 0.0001 | -0.83 |
| Turbidity | *Mychonastes* | 0.6013 | -0.13 |
| NO3 | *Mychonastes* | 0.1206 | -0.37 |
| PO4 | *Mychonastes* | 0.0757 | -0.42 |
| Temp_spatial | *Nannochloropsis* | 0.0097 | -0.58 |
| Salinity | *Nannochloropsis* | 0.3887 | -0.21 |
| Turbidity | *Nannochloropsis* | 0.6328 | 0.12 |
| NO3 | *Nannochloropsis* | 0.3708 | -0.22 |
| PO4 | *Nannochloropsis* | 0.0216 | -0.52 |
| Temp_spatial | *Minidiscus* | 0.7259 | 0.09 |
| Salinity | *Minidiscus* | 0.0001 | 0.77 |
| Turbidity | *Minidiscus* | 0.9488 | -0.02 |
| NO3 | *Minidiscus* | 0.1671 | 0.33 |
| PO4 | *Minidiscus* | 0.0096 | 0.58 |
| Temp_spatial | *Choricystis* | 0.6446 | -0.11 |
| Salinity | *Choricystis* | 0.2049 | 0.30 |
| Turbidity | *Choricystis* | 0.0517 | 0.45 |
| NO3 | *Choricystis* | 0.0068 | 0.60 |
| PO4 | *Choricystis* | 0.0150 | 0.55 |
| Temp_seasonal | Pico [cells/ mL] | < 0.0001 | 0.53 |
| Temp_spatial | Pico [cells/ mL] | < 0.0001 | 0.59 |
| Salinity | Pico [cells/ mL] | 0.2712 | 0.100 |
| Turbidity | Pico [cells/ mL] | 0.0291 | -0.2 |
| NO3 | Pico [cells/ mL] | < 0.0001 | -0.48 |
| PO4 | Pico [cells/ mL] | 0.1310 | -0.14 |
| Temp_seasonal | Pico [% of phyto] | 0.3255 | -0.09 |
| Temp_spatial | Pico [% of phyto] | 0.9793 | ~ 0 |
| Salinity | Pico [% of phyto] | 0.0397 | -0.19 |
| Turbidity | Pico [% of phyto] | < 0.0001 | 0.42 |
| NO3 | Pico [% of phyto] | < 0.0001 | 0.5 |
| PO4 | Pico [% of phyto] | 0.0010 | 0.29 |
| Temp_seasonal | Picoeuk [cells/ mL] | < 0.0001 | 0.47 |
| Temp_spatial | Picoeuk [cells/ mL] | < 0.0001 | 0.54 |
| Salinity | Picoeuk [cells/ mL] | 0.3039 | 0.09 |
| Turbidity | Picoeuk [cells/ mL] | 0.0254 | -0.20 |
| NO3 | Picoeuk [cells/ mL] | < 0.0001 | -0.50 |
| PO4 | Picoeuk [cells/ mL] | 0.0368 | -0.19 |
| Temp_seasonal | Picoeuk [% of phyto] | < 0.0001 | -0.48 |
| Temp_spatial | Picoeuk [% of phyto] | 0.0012 | -0.29 |
| Salinity | Picoeuk [% of phyto] | 0.2404 | -0.11 |
| Turbidity | Picoeuk [% of phyto] | < 0.0001 | 0.37 |
| NO3 | Picoeuk [% of phyto] | 0.0032 | 0.26 |
| PO4 | Picoeuk [% of phyto] | 0.5694 | -0.05 |
| Temp_seasonal | Picocyano [cells/ mL] | < 0.0001 | 0.70 |
| Temp_spatial | Picocyano [cells/ mL] | < 0.0001 | 0.63 |
| Salinity | Picocyano [cells/ mL] | 0.2698 | 0.10 |
| Turbidity | Picocyano [cells/ mL] | 0.0067 | -0.24 |
| NO3 | Picocyano [cells/ mL] | < 0.0001 | -0.41 |
| PO4 | Picocyano [cells/ mL] | 0.3389 | -0.09 |
| Temp_seasonal | Picocyano [% of phyto] | < 0.0001 | 0.42 |
| Temp_spatial | Picocyano [% of phyto] | 0.1551 | 0.13 |
| Salinity | Picocyano [% of phyto] | 0.2590 | -0.10 |
| Turbidity | Picocyano [% of phyto] | 0.0019 | 0.28 |
| NO3 | Picocyano [% of phyto] | < 0.0001 | 0.52 |
| PO4 | Picocyano [% of phyto] | < 0.0001 | 0.40 |

**Fig. S1:**
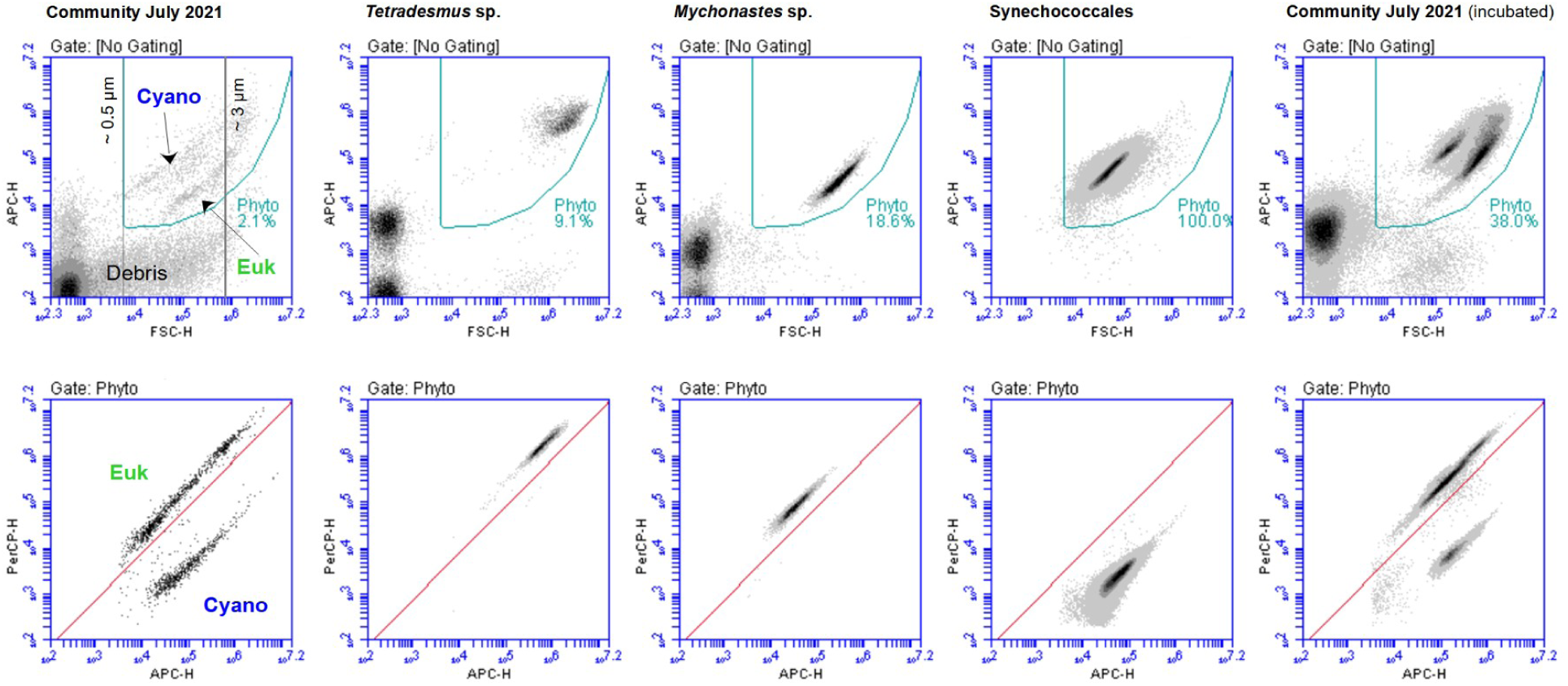
Cytogram examples of communities and single strains of phytoplankton isolated from the Elbe estuary. We used approximately 40 different samples (isolated taxa and laboratory incubated communities) from the Elbe estuary to determine the borders between phytoplankton and other suspended matter (e.g. bacteria, debris) and between eukaryotes and phycocyanin-rich cyanobacteria based on the flow cytometric properties. The gating of phytoplankton (phyto; upper panels) was based on forward scatter (FSC; ∼ size) and APC (allophycocyanin fluorescence; red fluorescence, filter 675 nm, laser 640 nm). Here we also set a lower size border of around 0.5 µm which was proxied in between the FSC values of 0.12 and 1 µm beads. The gating of eukaryotes (euk) and cyanobacteria (cyano) (lower panels) was done based on the ratio of APC and PerCP (peridinin-chlorophyll-protein complex fluorescence; red fluorescence, filter 670 nm, laser 488 nm). Lastly, we defined the picophytoplankton threshold based on 3 µm beads.

**Fig. S2:**
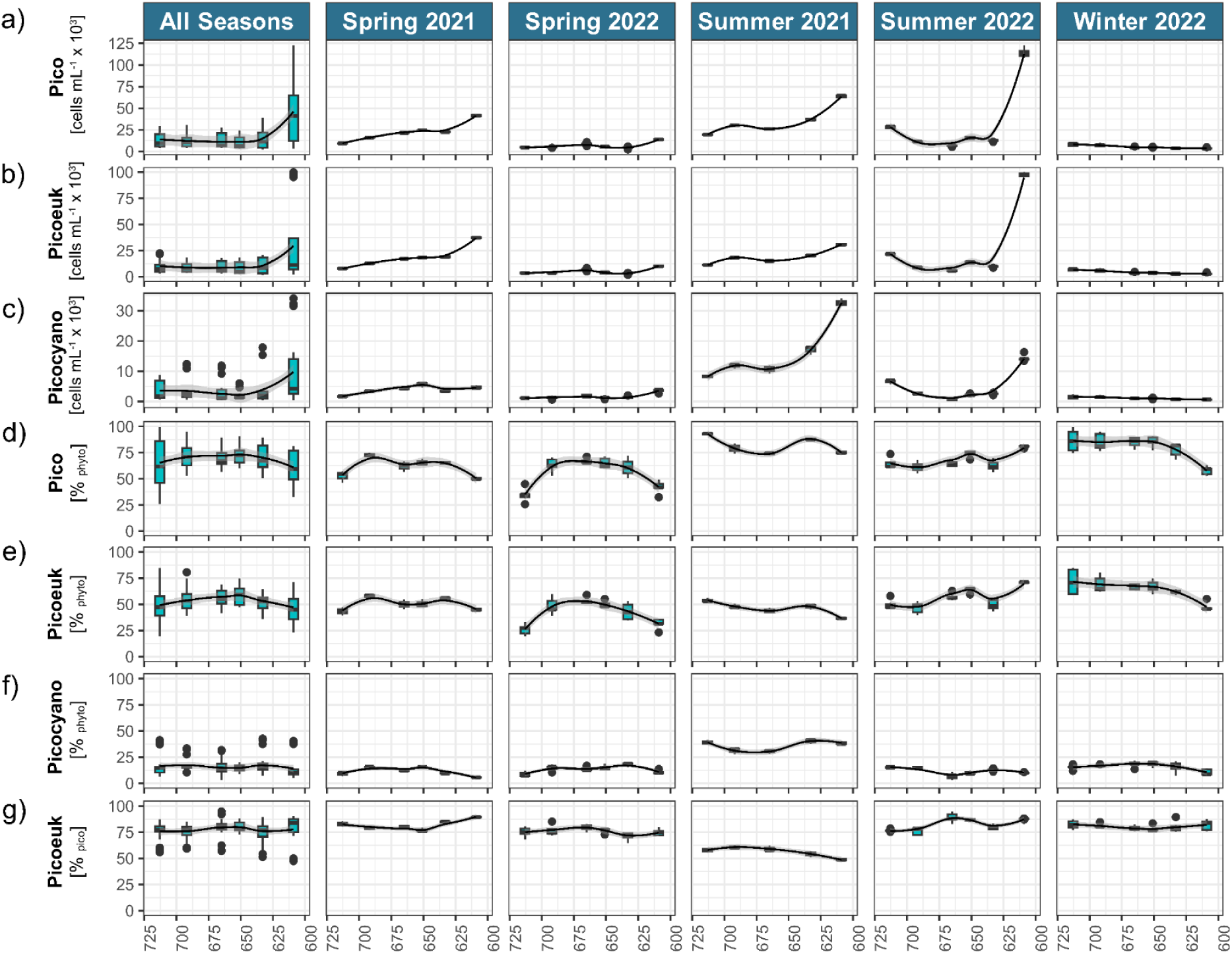
Spatial distribution of different picophytoplankton groups along different stations (stream km; fig. 1) and different seasons. Contributions of cyanobacteria to picophytoplankton can be obtained from c) (“picocyanobacteria = 1 - picoeukaryotes”). Regression lines were added with geom_smooth() from ggplot2 and the method “loess” as visual support. For clarity we use the following abbreviations: Pico = picophytoplankton, picoeuk = picoeukaryotes, phyto = phytoplankton. Note that 651 km was not sampled in July 2021.

**Fig. S3:**
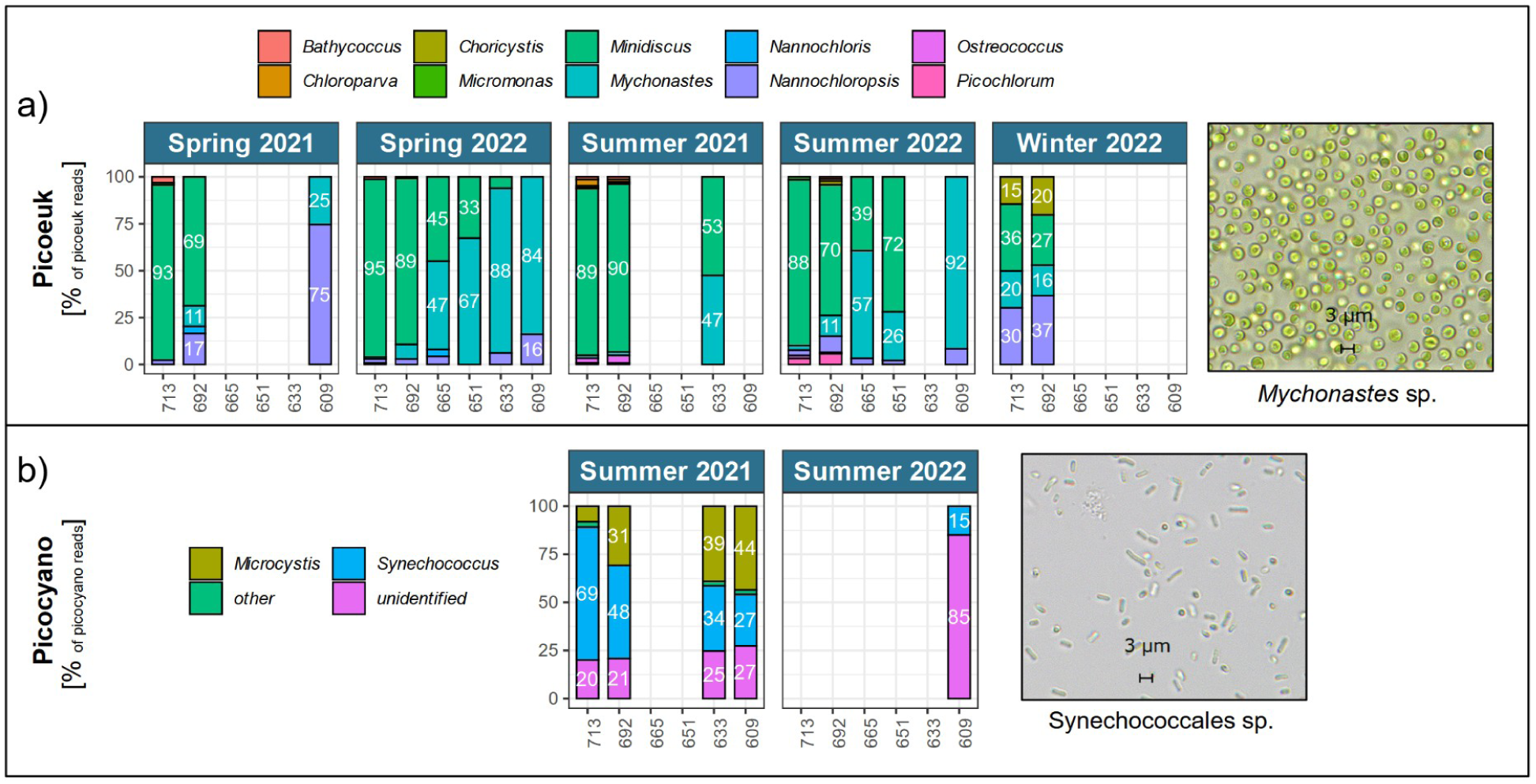
Spatial contributions of picoeukaryote genera to picoeukaryote 18S rRNA reads (a) and picocyanobacteria genera to picocyanobacteria 16S rRNA reads (b). Labels are shown where contribution was at least 10 %. Station is used as a factor (i.e. bars have the same distance independent of the actual location). Note that in each a) and b) we selected different taxa that *can* be or are *usually* understood as picophytoplankton (< 3 µm). Cells of these taxa however *can* sometimes be larger than 3 µm or form colonies and some unidentified cyanobacteria might not be < 3 µm. Moreover, the number of reads can depend on cell size (Godhe et al., 2008). Hence, the selection in this plot does not necessarily accurately represent the picophytoplankton obtained from flow cytometry, where the definition is based on their actual cell size and abundance (see also fig. 1b). Where bars are missing - or in case of picocyanobacteria even complete panels - metabarcoding was either not carried out (651 - 665 km in 2021) or data was removed due to low number of picocyanobacteria or picoeukaryotes reads (< 100) (all other cases). Photos show examples of picophytoplankton strains isolated from the Elbe estuary (Martens et al., 2024a). For clarity we use the following abbreviations: Picocyano = picocyanobacteria, picoeuk = picoeukaroytes.

**Fig. S4:**
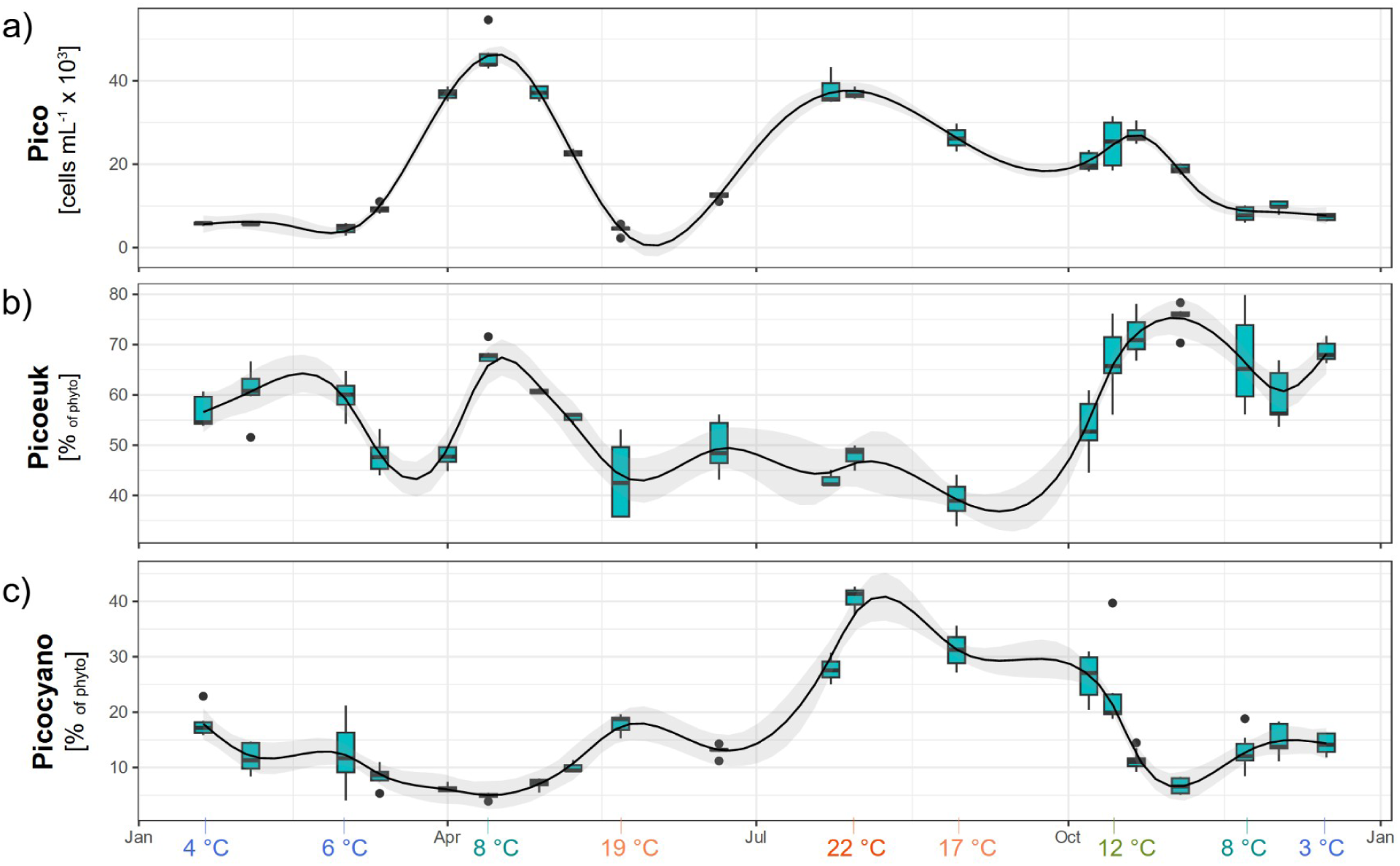
Seasonal distribution of different picophytoplankton groups in the area around Hamburg (approx. 623 - 633 km). Horizontal scales show the sampling date independent of the year, i.e. day of the month. Data was merged when sampling was carried out < 5 days apart. Contributions of cyanobacteria to picophytoplankton can be obtained from the picoeukaryotes contribution in c) (“picocyanobacteria = 1 - picoeukaryotes”). Regression lines were added with geom_smooth() from ggplot2 and the method “gam” (see also tab. S3). For clarity we use the following abbreviations: Pico = picophytoplankton, picoeuk = eukaryotes, phyto = phytoplankton. On the bottom we show the temperatures at certain time points (see further details in fig. S6).

**Fig. S5:**
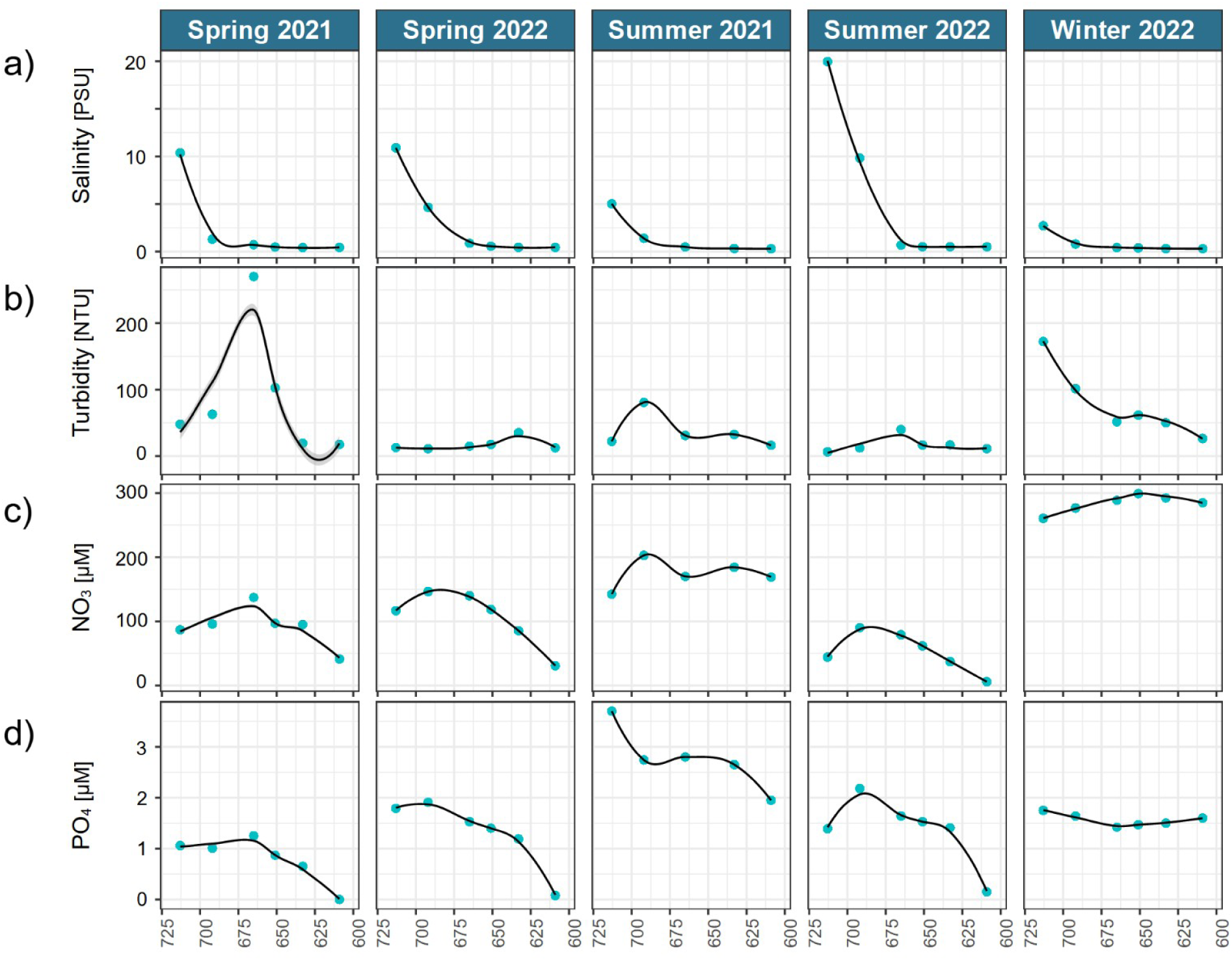
Abiotic conditions at the different stations and seasons of the spatial dataset. Regression lines added with geom_smooth() from ggplot2 and the method “loess” as visual support. Note that one missing value of turbidity on 2021-07-29 (summer 2021) at 609 km was replaced by a value from 2021- 07-26 at 609 km.

**Fig. S6:**
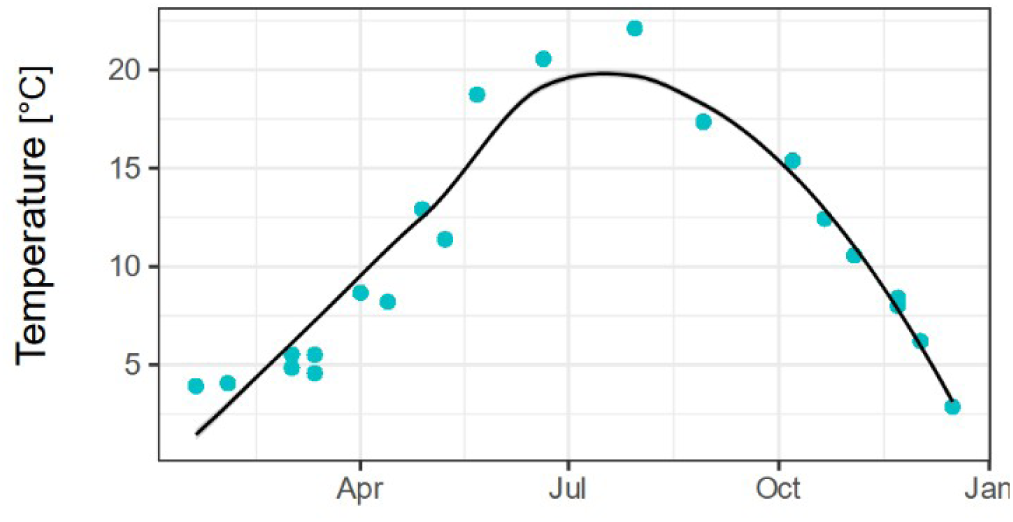
Water temperature along seasons in the seasonal dataset. Regression line added with geom_smooth() from ggplot2 and the method “loess” as visual support.

**Fig. S7:**
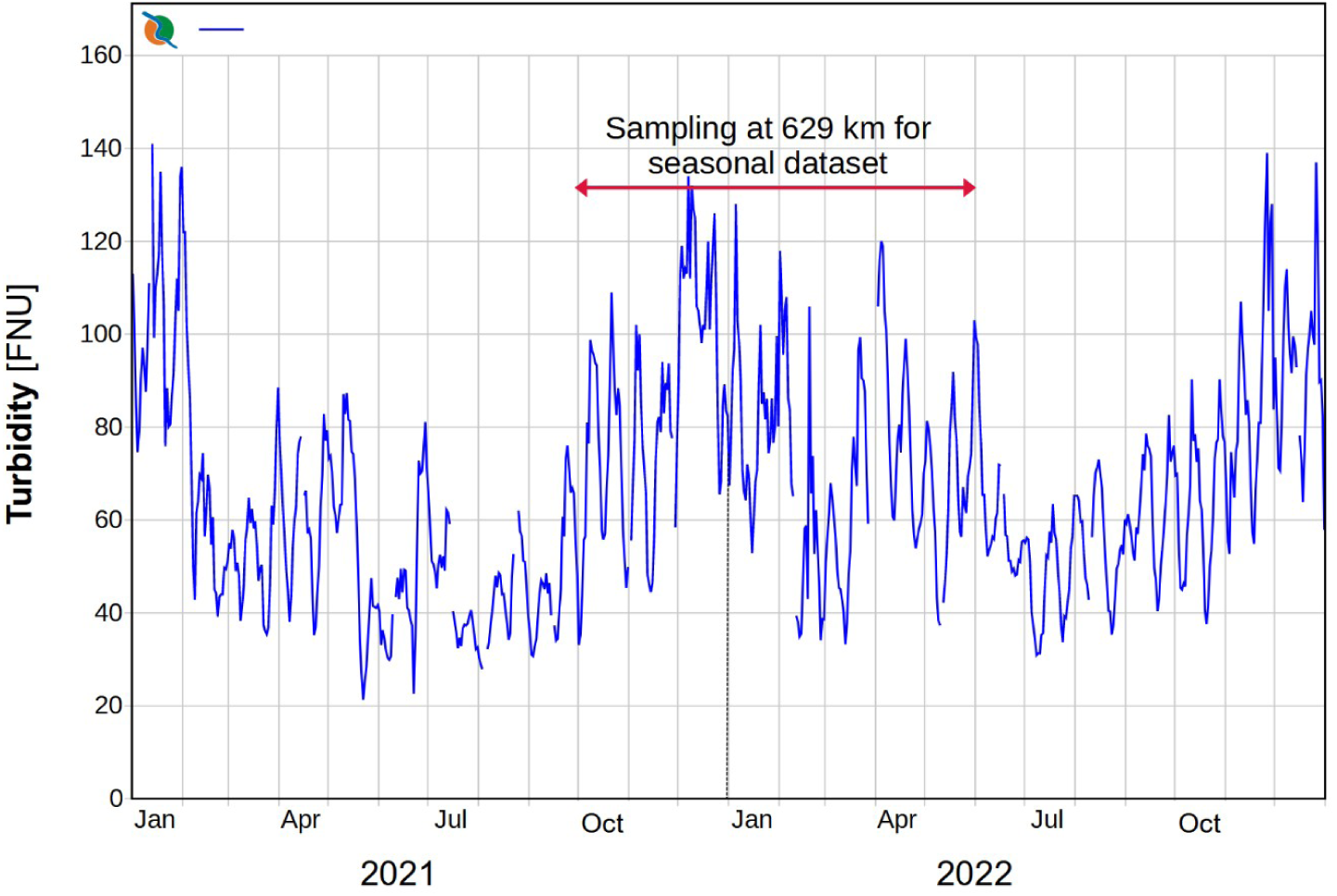
Average daily turbidity (FNU) at the station Seemannshöft (approx. 629 km) in the Elbe estuary based on data from the FGG Elbe database (FGG Elbe, 2024) from 2021 - 2022. Figure was obtained directly from the FGG database, but modified by removing German texts.

## Notes

### Competing Interest Statement

The authors have declared no competing interest.

